# Dynamics of postnatal bone development and epiphyseal synostosis in the caprine autopod

**DOI:** 10.1101/2024.12.26.630423

**Authors:** Christopher J. Panebianco, Maha Essaidi, Elijah Barnes, Ashley Williams, Karin Vancíková, Margot C. Labberté, Pieter Brama, Niamh C. Nowlan, Joel D. Boerckel

**Author notes:** Corresponding authors: Niamh C. Nowlan University College Dublin UCD School of Mechanical and Materials Engineering Belfied, Dublin 4, Ireland (Email) Joel D. Boerckel University of Pennsylvania Department of Orthopaedic Research Department of Bioengineering 376A Stemmler Hall 3450 Hamilton Walk Philadelphia, PA 19104-6081 (Email).

## Abstract

Bones develop to structurally balance strength and mobility. Bone developmental dynamics are influenced by whether an animal is ambulatory at birth (*i.e.,* precocial). Precocial species, such as goats, develop advanced skeletal maturity in utero, making them useful models for studying the dynamics of bone formation under mechanical load. Here, we used microcomputed tomography and histology to characterize postnatal bone development in the autopod of the caprine lower forelimb. The caprine autopod features two toes, fused by metacarpal synostosis (*i.e.*, bone fusion) prior to birth. Our analysis focused on the phalanges 1 (P1) and metacarpals of the goat autopod from birth through adulthood (3.5 years). P1 cortical bone densified rapidly after birth (half-life using one-phase exponential decay model (τ_1/2_ = 1.6 ± 0.4 months), but the P1 cortical thickness increased continually through adulthood (τ_1/2_ = 7.2 ± 2.7 mo). Upon normalization by body mass, the normalized polar moment of inertia of P1 cortical bone was constant over time, suggestive of structural load adaptation. P1 trabecular bone increased in trabecular number (τ_1/2_ = 6.7 ± 2.8 mo) and thickness (τ_1/2_ = 6.6 ± 2.0 mo) until skeletal maturity, while metacarpal trabeculae grew primarily through trabecular thickening (τ_1/2_ = 7.9 ± 2.2 mo). Unlike prenatal fusion of the metacarpal diaphysis, synostosis of the epiphyses occurred postnatally, prior to growth plate closure, through a unique fibrocartilaginous endochondral ossification. These findings implicate ambulatory loading in postnatal bone development of precocial goats and identify a novel postnatal synostosis event in the caprine metacarpal epiphysis.

## 1.0 Introduction

The skeleton continuously remodels throughout mammalian growth and development, creating a structure optimized for high strength and low weight. While we have long known that mature bone adapts and remodels based on loading (Wolff 1893; Frost 1994), it has been difficult to decouple early bone development programs from adaptation to ambulatory loading. As precocial species, goats are ambulatory at birth, providing continuous loading during skeletal growth. Here, we mapped the developmental dynamics of the autopod of the caprine lower forelimb from birth through adulthood, focusing on the phalanges 1 (P1) and metacarpal bones in the autopod.

An animal’s ability to ambulate at birth influences the dynamics of postnatal skeletal development. Humans, mice, and rats are examples of altricial species, meaning they are non-ambulatory at birth. Altricial species initiate primary ossification in utero, under conditions of dynamic fetal movement (Nowlan 2015; Beall et al. 2007; Kozhemyakina et al. 2015; Collins et al. 2024), then the majority of secondary ossification centers initiate after birth (Kozhemyakina et al. 2015; Breeland et al. 2024; Gardner and Gray 1970). Unlike altricial species, precocial species such as goats, horses, and sheep, ambulate from birth through adulthood (**Fig. 1**) and undergo primary and secondary ossification in utero as an anticipatory mechanism to prepare for future loading (Gorissen et al. 2016; Skedros et al. 2004; Skedros et al. 2007). It has been hypothesized that this early secondary ossification evolved to support the physis to withstand high mechanical loading shortly after birth (Xie et al. 2020). Though previous research has described gestational bone development of precocial species, few studies have investigated their postnatal development. Understanding postnatal developmental dynamics in precocial species may inform how mechanical cues influence skeletal morphogenesis.

**Figure 1.**
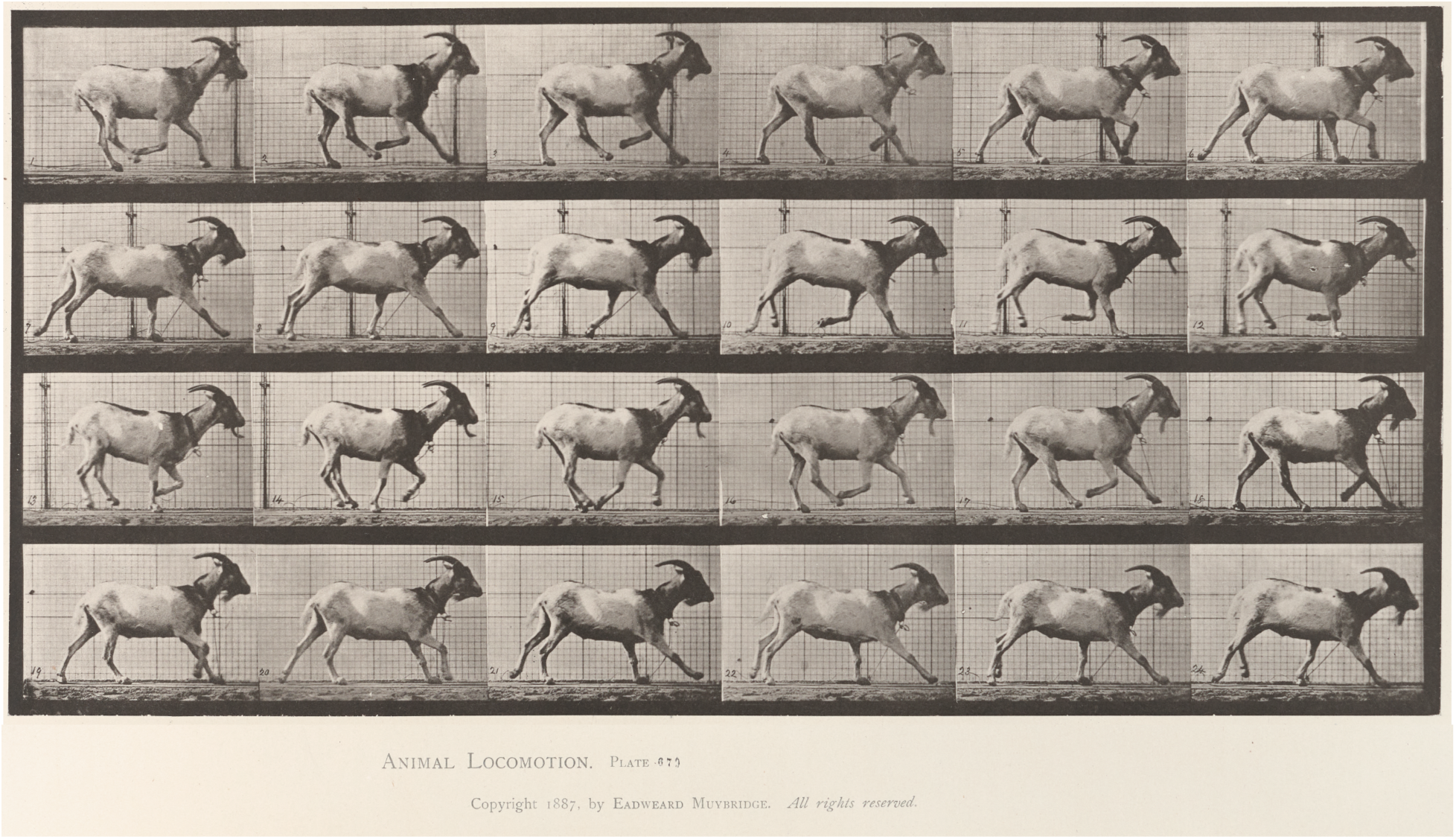
Goats are precocial animals that ambulate from birth through adulthood. Reproduction of 1887 Animal Locomotion Plate 676, captured by Eadweard Muybridge at the University of Pennsylvania. Plates provided by the University of Pennsylvania University Archives and Records Center.

Diaphyseal synostosis has evolved in a variety of species, likely to provide structural resistance to high bending forces. For example, the jerboa, a bipedal rodent, evolved fused metatarsal bones that reduce peak bone stresses and provide a factor-of-safety to the hindlimb during bipedal ambulation (Villacís Núñez et al. 2022). This evolutionary advantage contributes to the jerboa’s ability to jump ten-times its hip-height and withstand peak ground-reaction forces that are five-times its body weight (Moore et al. 2017; Biewener 1989). As altricial species with limited hindlimb loading immediately after birth, jerboa metatarsal synostosis occurs postnatally (Gutierrez et al. 2019; Cooper et al. 2014). Diaphyseal synostosis is also observed in some precocial species. For example, each goat forelimb has two independent metacarpal rudiments embryonically, but these rudiments undergo diaphyseal synostosis in utero to form one metacarpal bone at birth. As distal elements (*e.g.,* autopod) mature later in development than the more proximal zeugopod and stylopod, evolutionary reasoning suggests that the ungulate diaphyseal synostosis initiates in utero to provide increased strength and enable locomotion at birth (Leinders and Sondaar 1973). It remains unknown whether these fusion events continue through postnatal development. These findings may be useful for understanding how biomechanical pressures guide adaptation in precocial species.

Here, we characterize the dynamics of postnatal bone development and epiphyseal synostosis in the caprine autopod from birth through adulthood. P1 cortical bone rapidly densified, then became continually thicker until skeletal maturity, with structural distribution determined by ambulatory loading. Simultaneously, P1 trabecular bone increased in trabecular number and thickness until skeletal maturity, while metacarpal trabecular number was constant and the network grew through trabecular thickening. Metacarpal synostosis began *in utero*; however, the epiphyses did not begin to fuse until after 1.5 months postnatally. Histological analyses indicated that this epiphyseal synostosis occurred through fibrocartilaginous endochondral ossification, prior to growth plate closure. Together, these data provide new insights into precocial bone development under ambulatory loading.

## 2.0 Materials and methods

### 2.1 Study design

We performed bone morphometry on the autopods of the lower forelimbs of 44 goats at the following ages: 3 days (3D, N=6 male, N=3 female), 1.5 months (1.5M, N=3 male, N=3 female), 3M (N=1 male, N=4 female), 6M (N=6 female), 9M (N=3 male, N=3 female), 12M (N=3 male, N=3 female), and 3.5 years (3.5Y, N=6 female). Each animal was considered an experimental unit and goats were assigned to groups in cohorts. Goats were continually assessed by a veterinarian, and only animals in good physical health were included in the study. At the specified timepoints, animals were euthanized and the phalanges 1 (P1) and metacarpal bones were harvested from autopods of the lower forelimbs for microcomputed tomography (µCT) and histological analyses.

### 2.2 Animal husbandry and care

This animal experiment complied with the ARRIVE guidelines. Ethical evaluation and approval were provided by the Animal Research Ethics Committee (AREC-21-22) of University College Dublin (UCD) and the Lyons UCD Research Farm Animal Welfare Board (Health, Husbandry and Monitoring plans: 201,907). Animals were selected at predetermined age points from the Translational Research Unit (TRU) goat herd at University College Dublin Lyons Research Farm (Breeding license AREC-22-408929). In principle all animals genetically originated from the same high health closed goat herd of our sole supplier.

The TRU goat herds were kept under normal goat husbandry conditions. Herds had loose housing and pasture access when weather conditions were appropriate. Daily health checks of the herd were performed by appropriately trained husbandry staff, with weather conditions and barn climate parameters documented on the daily health check sheets. Animals were reared with their respective mothers when available or at a lambing bar with milk replacer to suckle milk until weaning at approximately 60-90 days (depending on body condition score and weight). Ad libitum concentrates and hay/silage were provided from approximately 14 days of age. After goats were 3 months, hay/silage remained ad libitum, but concentrates were provided based on their daily requirements and weight gain to prevent overfeeding. Veterinary staff of the TRU performed weekly monitoring health checks on the goat herds and were available 24/7 in case a veterinary consultation is required.

When animals reached their predetermined age endpoint, a normal general health check and orthopaedic assessment was conducted. Only animals that met these requirements were included in the study. Included animals underwent humane euthanasia under sedation with a barbiturate overdose. Following euthanasia, the P1 and metacarpal bones were harvested for microcomputed tomography (µCT) and histological analyses. Samples were fixed using 10% neutral buffered formalin (NBF) for 3 days, then 5% NBF for 3 weeks. Fixed samples were washed with 1X phosphate buffered saline (PBS), then frozen at −20°C in a solution of 30% sucrose (w/v) in 1X PBS prior to analyses.

### 2.3 Microcomputed tomography (µCT)

The night before scanning, frozen samples were thawed overnight at 4°C. Thawed samples were wrapped in PBS-soaked gauze and scanned using a SCANCO µCT 45 desktop microCT scanner (SCANCO Medical, Wangen-Brüttisellen, Switzerland). X-ray images were acquired using x-ray intensity of 145 µA, energy of 55 kVp, integration time of 400 ms, and resolution of 10.4 µm. Reconstructed scans were semi-automatically contoured and analyzed using the SCANCO microCT image analysis interface (SCANCO Medical) according to morphometric standards (Bouxsein et al. 2010).

For P1 bones, approximately 200 slices of the midshaft cortical region were contoured to measure cortical tissue cross-sectional area (Tt.Ar), cortical bone area (Ct.Ar), cortical area fraction (Ct.Ar/Tt.Ar), cortical thickeness (Ct.Th), tissue mineral density (TMD), and polar moment of inertia (*J*). Additionally, the entire P1 distal trabecular region was contoured to measure trabecular bone volume (BV/TV), trabecular number (Tb.N), trabecular thickness (Tb.Th), and trabecular separation (Tb.Sp). For each goat, the cortical and trabecular outcome measurements were averaged for the medial and lateral P1 bones, so each goat was an experimental unit.

For metacarpal bones, BV/TV, Tb.N, Tb.Th, and Tb.Sp were calculated for the combined distal trabecular network in both epiphyses. Additionally, the maximum epiphyseal fusion length was calculated for each metacarpal bones using the mid-sagittal virtual slice numbers for the initiation of epiphyseal synostosis and the physis.

### 2.4 Histology

Representative P1 and metacarpal samples (N=3 per timepoint) were used for paraffin histology. P1 samples were decalcified for 3.5 weeks using Formical-4 Decalcifier (StatLab Medical Products, McKinney, Texas, USA), with weekly reagent changes. Metacarpal samples were decalcified for 4 weeks using Formical-4, then an additional 2 weeks using Formical-2000 (StatLab Medical Products), with weekly reagent changes. Decalcified samples were trimmed using a Tissue-Tek® Accu-Edge® Trimming Knife (Electron Microscope Sciences, Hatfield, PA, USA) and processed for paraffin histology using a Thermo Scientific Excelsior AS Tissue Processor (Thermo Fisher Scientific, Waltham, MA). Processed samples were paraffin-embedded and sectioned to 10 µm using a Leica RM 2030 Microtome (Leica, Bala Cynwyd, PA, USA). Sections were stained for safranin-O/fast green using standard protocols. Stained sections were imaged using a ZEISS Axio Scan.Z1 Slide Scanner (ZEISS, Oberkochen, Germany).

### 2.5 Statistical analysis

The “pwr2” package in R (Indianapolis, IN, USA) was used to conduct a power analysis for large effect sizes (**α**=0.05, **β**=0.20, f = 0.4-0.6). GraphPad Prism software Version 10.3.1 (GraphPad Software, San Diego, CA, USA) was used to conduct all statistical analyses. A one-way analysis of variance (ANOVA) with Tukey’s multiple comparison test was used to determine significant differences in all outcome measurements over developmental time. Prism was also used to fit all outcome measurements with a one-phase exponential decay model 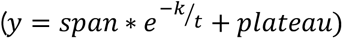. Graphical results are reported as a one-phase decay model with the goodness of fit (R^2^) and half-life (τ_1/2_). τ_1/2_ is the amount of developmental time, in months, for each outcome to achieve 50% of its maximum value.

## 3.0 Results

### 3.1 Phalanges 1 (P1) postnatal development dynamics

Phalanges 1 (P1) bones showed significant increases in all outcomes for cortical bone morphometry over postnatal development (**Fig. 2**). To better understand the dynamics of caprine P1 development over time, each outcome measurement was fit with a one-phase exponential decay model. Curves fit well for all outcomes with coefficients of determination (R^2^) values between 0.62 - 0.91 (**Fig. 2B-G**). By extracting the half-life values (τ_1/2_) from these curves, we could determine the characteristic time scale for each outcome. Outcomes with lower τ_1/2_ values achieved 50% of their maximum value earlier in developmental time than outcomes with higher τ_1/2_ values. Thus, outcomes with lower τ_1/2_ values achieved their maximum value faster and the outcome plateaued earlier in developmental time than outcomes with higher τ_1/2_ values. Cortical area fraction (Ct.Ar/Tt.Ar, τ_1/2_ = 1.0 ± 0.3 mo, **Fig. 2D**), a measure of cortical porosity, and tissue mineral density (TMD, τ_1/2_ = 1.6 ± 0.4 mo, **Fig. 2E**) plateaued most rapidly. Total cross-sectional area (Tt.Ar, τ_1/2_ = 4.8 ± 1.7 mo, **Fig. 2B**), cortical bone area (Ct.Ar, τ_1/2_ = 6.0 ± 2.6 mo, **Fig. 2C**), cortical thickness (Ct.Th, τ_1/2_ = 7.2 ± 2.7 mo, **Fig. 2F**), and polar moment of inertia (*J*, τ_1/2_ = 4.3 ± 1.9 mo, **Fig. 2G**) plateaued less rapidly and continued to increase through 12 months. These results indicate that P1 bones undergo early cortical condensation followed by progressive cortical thickening into adulthood.

**Figure 2.**
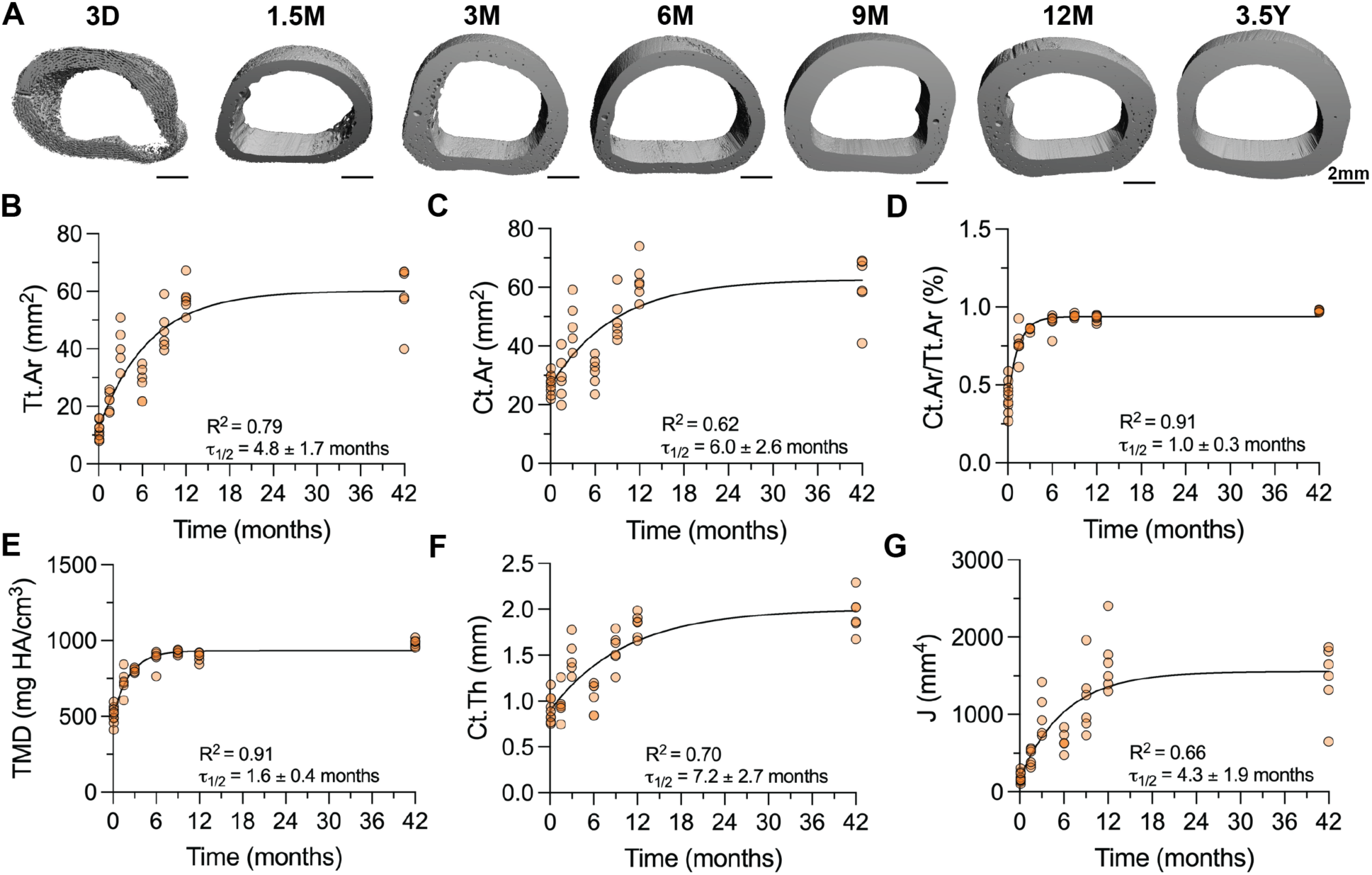
Phalanges 1 (P1) cortices exhibit rapid condensation and continuous expansion until adulthood. **(A)** Transverse views of 3D microcomputed tomography (microCT) reconstructions of P1 cortical mid-shaft. Scale bars = 2 mm. **(B)** Cortical tissue cross-sectional area (Tt.Ar), **(C)** Cortical bone area (Ct.Ar), **(D)** Cortical area fraction (Ct.Ar/Tt.Ar), **(E)** Tissue mineral density (TMD), **(F)** Cortical thickness (Ct.Th), and **(G)** Polar moment of inertia (J) throughout developmental time, fit with a one-phase decay model. Goodness of fit (R^2^) and half-life (τ_1/2_) are reported for each graph. D = days of age, M = months of age, Y = years of age.

P1 distal trabecular microarchitecture also showed significant changes in all outcomes over postnatal development (**Fig. 3**). Trabecular number (Tb.N, τ_1/2_ = 6.7 ± 2.8 mo, **Fig. 3B**) and trabecular thickness (Tb.Th, τ_1/2_ = 6.6 ± 2.0 mo, **Fig. 3C**) increased continuously through 12 months. As a result of this increased formation of new trabecular networks and growth of existing trabecular networks, the bone volume fraction (BV/TV, τ_1/2_ = 3.3 ± 1.0 mo, **Fig. 3D**) increased and the trabecular separation (Tb.Sp, τ_1/2_ = 7.9 ± 3.2 mo, **Fig. 3E**) decreased continuously through 12 months. Simultaneously, the existing trabeculae were increasing in mineralization, as shown by the continuous changes in TMD (τ_1/2_ = 4.2 ± 1.7 mo, **Fig. 3F**). Curves fit well for all outcomes with R^2^ values between 0.63 - 0.85 (**Fig. 3B-F**). These data suggest that P1 postnatal cancellous bone formation is characterized by both the formation of new trabecular networks and the growth of existing trabecular networks.

**Figure 3.**
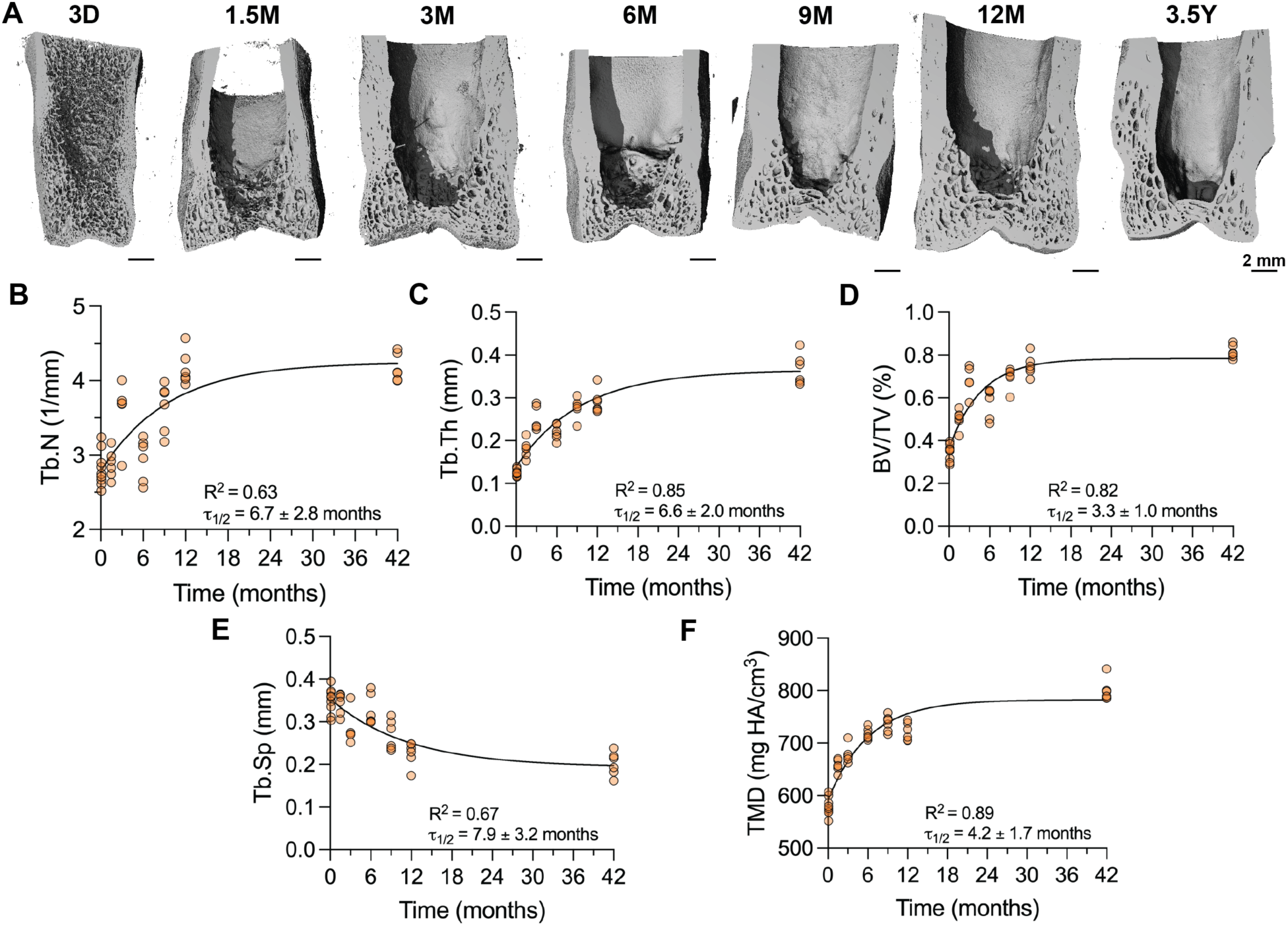
Distal P1 bones grow by formation of new trabeculae and expansion of existing trabeculae. **(A)** Transverse views of 3D microCT reconstructions of P1 bones with a virtual slice through the mid-coronal plane using microCT. Scale bars = 2 mm. **(B)** Trabecular number (Tb.N), **(C)** Trabecular thickness (Tb.Th), **(D)** Bone volume fraction (BV/TV), **(E)** Trabecular spacing (Tb.Sp), and **(F)** Tissue mineral density (TMD) throughout developmental time, fit with a one-phase decay model. Goodness of fit (R^2^) and half-life (τ_1/2_) are reported for each graph. D = days of age, M = months of age, Y = years of age.

P1 bone structure formed concurrently with precocial ambulatory weight-bearing. Animals increased continuously in mass from birth through adulthood (τ_1/2_ = 5.1 ± 1.2 mo, **Fig. 4A**). To determine how bone morphometry adapted with loading, we normalized cortical and trabecular outcomes by animal body mass. Body mass-normalized Ct.Ar (τ_1/2_ = 0.9 ± 0.4 mo, **Fig. 4B**), cortical TMD (τ_1/2_ = 1.4 ± 0.5 mo, **Fig. 4C**), BV/TV (τ_1/2_ = 1.5 ± 0.5 mo, **Fig. 4E**), and Tb.Th (τ_1/2_ = 1.3 ± 0.5 mo, **Fig. 4F**) all plateaued rapidly. In contrast, the body mass-normalized polar moment of inertia (*J/*mass) was constant over time (**Fig. 4D**). These data suggest that while bone accrual was dominated by animal growth rather than load-adaptation, the morphodynamics of cross-sectional distribution, indicative of structure resistance to bending and torsional loading, were adaptive to ambulatory loads.

**Figure 4.**
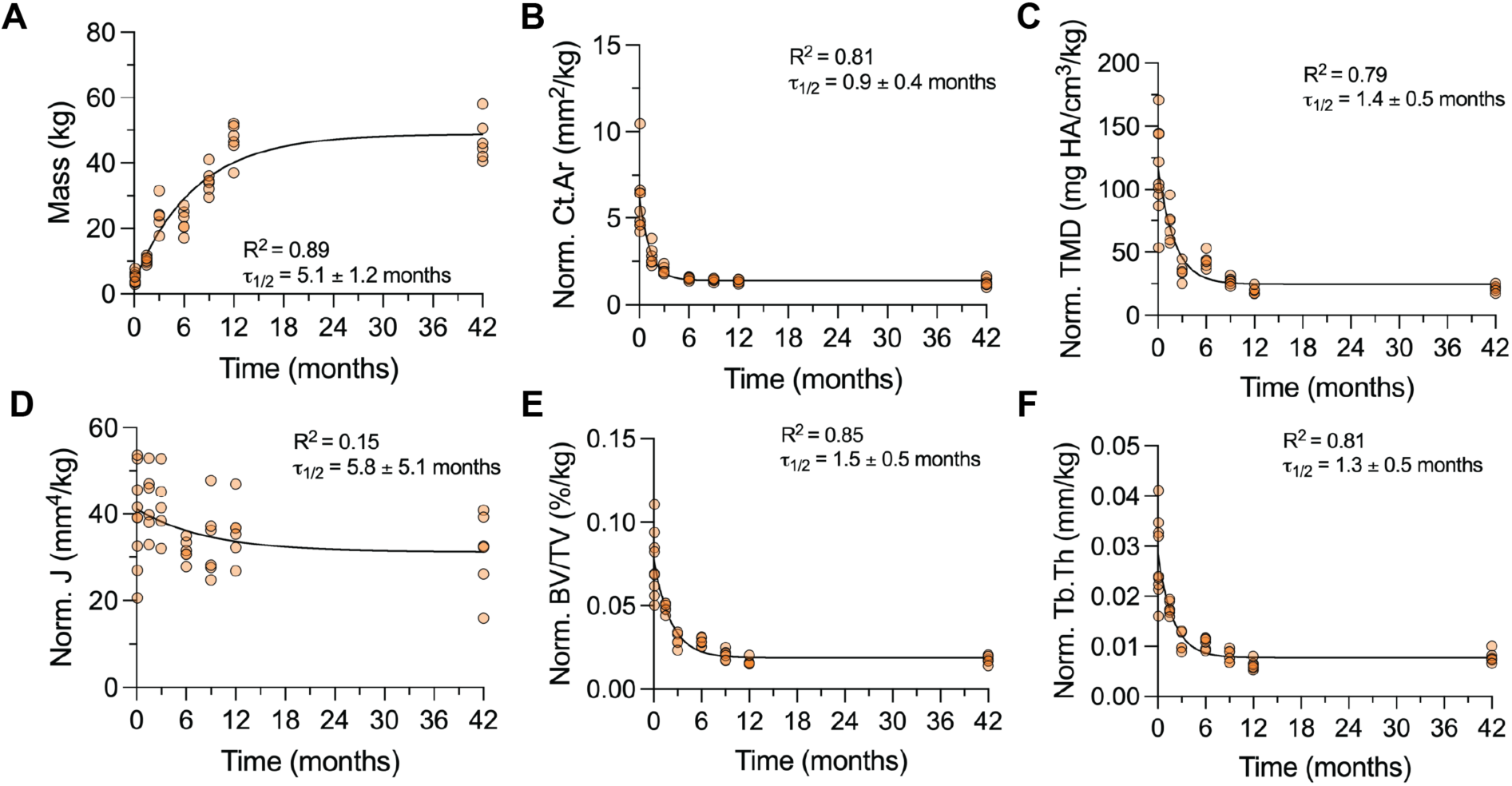
Morphodynamics of P1 cross-sectional distribution are adaptive to ambulatory loads. **(A)** Animal mass throughout developmental time. Body mass normalized **(B)** Cortical bone area (Ct.Ar), **(C)** Cortical tissue mineral density (TMD), **(D)** Polar moment of inertia (J), **(E)** Trabecular bone volume fraction (BV/TV), and **(F)** Trabecular thickness (Tb.Th). Prefix “Norm” indicates outcome is normalized to animal mass. Data graphed as individual data points vs. time fit with a one-phase decay model. Goodness of fit (R^2^) and half-life (τ_1/2_) are reported for each graph.

### 3.2 Metacarpal postnatal development dynamics

Synostosis of the metaphyseal diaphyses was uniformly present in neonates, indicating fusion during fetal development (**Fig. 5A**). Cortical fusion was only observed in the diaphyseal region of the metacarpals and preserved the structure of two distinct distal physes. Additionally, metacarpal bones maintained distinct lateral and medial trabecular networks in the diaphysis and metaphysis.

**Figure 5.**
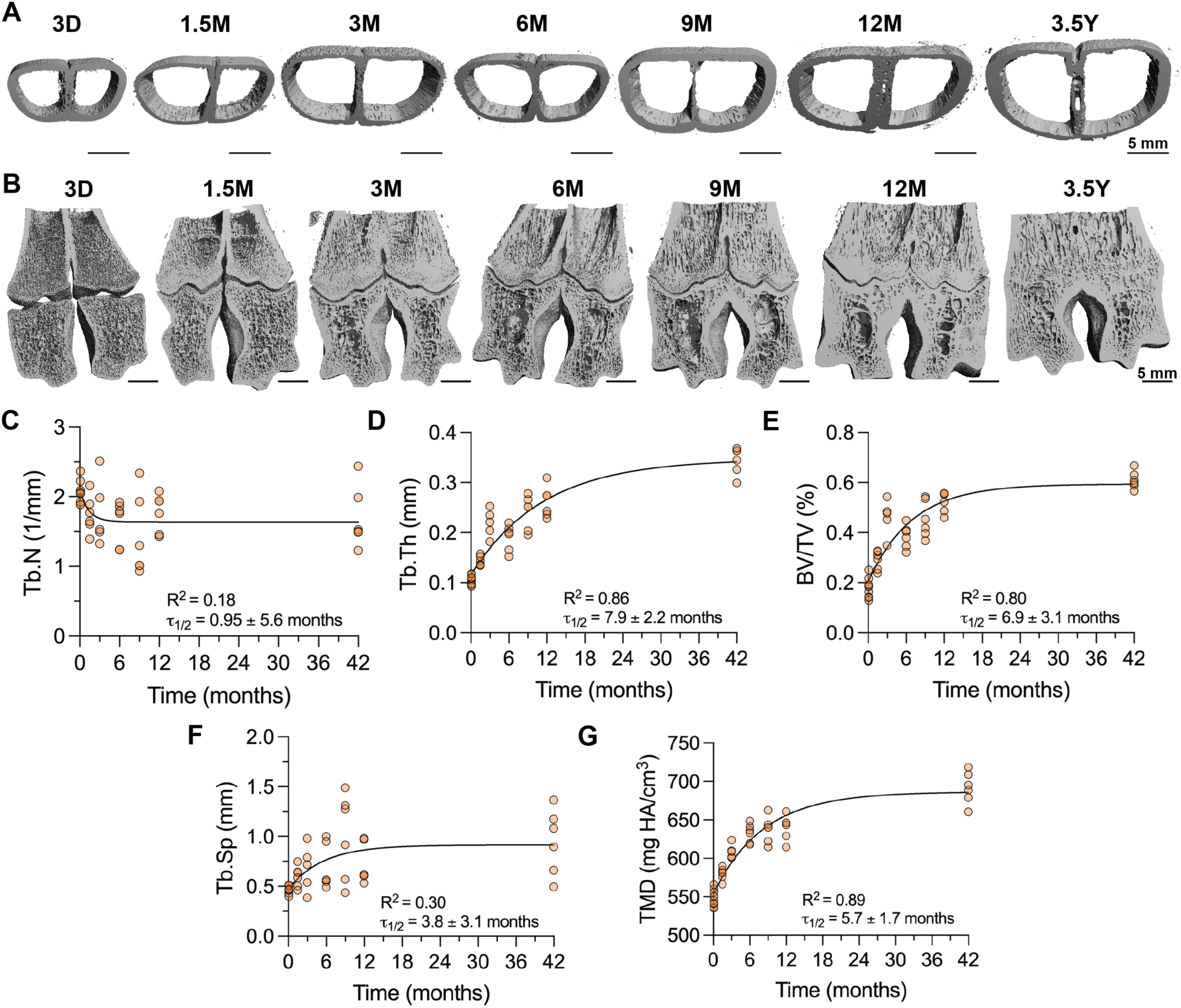
Metacarpal cortices fuse prior to birth and distal metacarpal trabecular bone grows through trabecular thickness expansion. (A) 3D microCT reconstructions of cortices with a virtual slice through the transverse plane, showing cortical fusion prior to birth. Scale bars = 5 mm. (B) 3D microCT reconstructions of metacarpal bones with a virtual slice through the mid-coronal plane. Scale bars = 5 mm. (C) Trabecular number (Tb.N), (D) Trabecular thickness (Tb.Th), (E) Bone volume fraction (BV/TV), (F) Trabecular spacing (Tb.Sp), and (G) Tissue mineral density (TMD) throughout developmental time, fit with a one-phase decay model. Goodness of fit (R^2^) and half-life (τ_1/2_) are reported for each graph. D = days of age, M = months of age, Y = years of age.

Bone formation in the epiphyseal metacarpal trabecular bone occurred by increasing trabecular thickness, rather than increasing trabecular number (**Fig. 5B-G**). Tb.N changed minimally during postnatal metacarpal development (**Fig. 5C**), while Tb.Th (τ_1/2_ = 7.9 ± 2.2 mo, **Fig. 5D**) increased continuously over 3.5 years, suggesting bone accrual on existing trabecular networks over time. Simultaneously, the trabecular bone matrix increased in mineral density, TMD (τ_1/2_ = 5.7 ± 1.7 mo, **Fig. 5F**). Despite limited changes in Tb.N, the continuously increasing Tb.Th was sufficient to increase trabecular BV/TV, but not Tb.Sp, over postnatal development (τ_1/2_ = 6.9 ± 3.1 mo, **Fig. 5E-F**). These findings suggest that the development of the epiphyseal metacarpal trabecular microarchitecture may feature mineral apposition on trabecular networks present at birth, rather than de novo trabecular formation.

### 3.3 Epiphyseal bone fusion dynamics in the metacarpal bones

Diaphyseal synostosis of the metacarpal bones occurred prenatally, but epiphyseal synostosis occurred postnatally. Diaphyseal synostosis could be observed as early as 3D (**Fig. 5A & 6A**), suggesting this fusion event occurred prior to birth. In contrast, synostosis of the epiphyses distal to the growth plate did not initiate until after 3 months (**Fig. 6A & B**). We quantified the extent of epiphyseal synostosis as the fusion length, which increased continuously between 3 months and 12 months. The distal metacarpal physis closed between 12 months and 3.5 years; thus, it was not possible to measure fusion length past 12 months. At 3.5 years, the metacarpal bone remodeled the growth plate into a mature network of cancellous bone.

**Figure 6.**
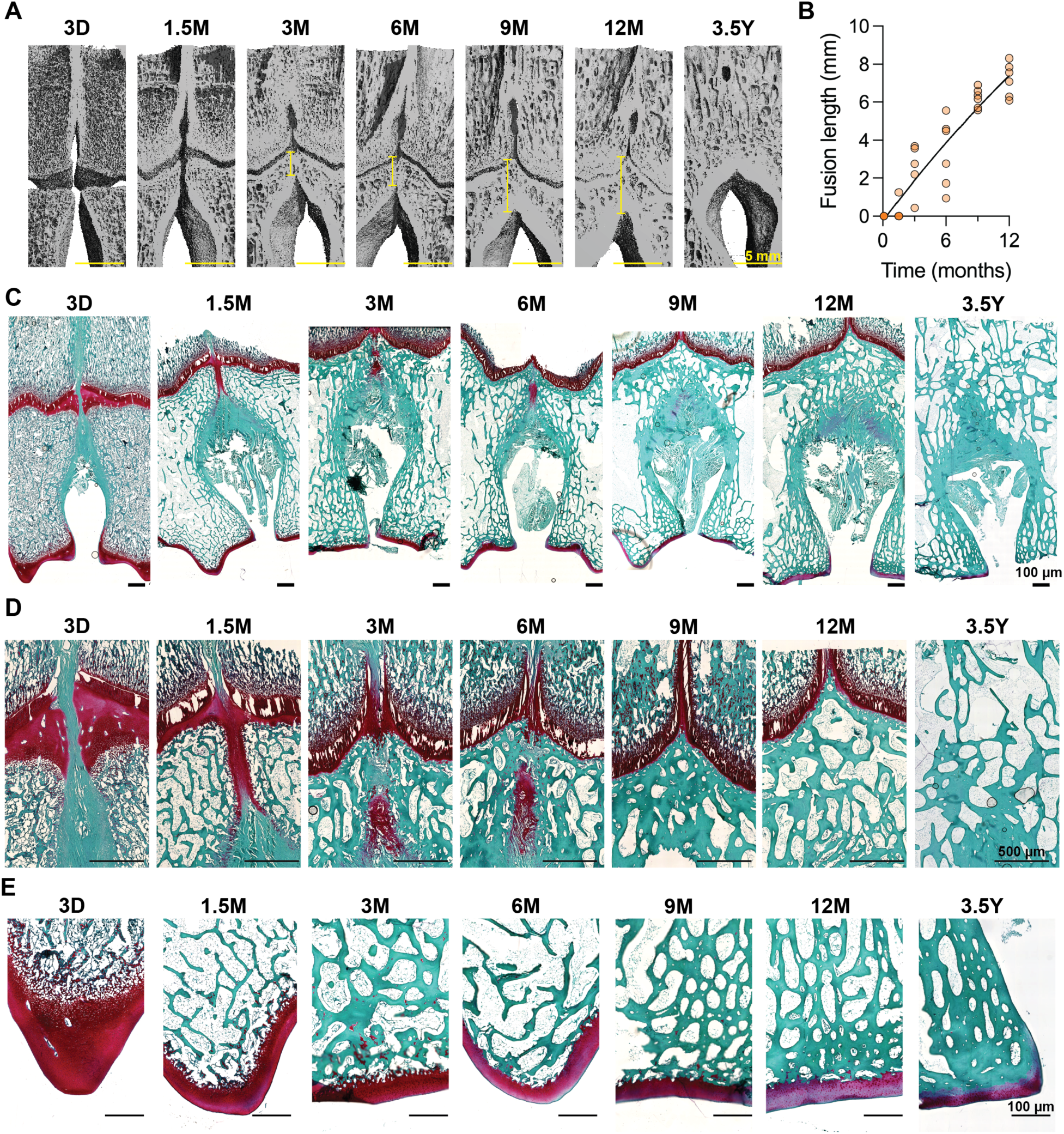
Metacarpal epiphyses are separated by a non-cartilaginous fibrous template at birth, but fuse by fibrocartilaginous endochondral ossification. (A) 3D microCT reconstructions of metacarpal bones with a virtual slice through the mid-coronal plane. Yellow line indicates extent of epiphyseal fusion (i.e., “fusion length”). Scale. bars = 5 mm. (B) Quantification of the epiphyseal fusion length throughout developmental time. (C) Safranin o/fast green histological micrographs of metacarpal bones cut through the mid-coronal plane. Scale bars = 100 µm. (D) High-magnification micrographs of the distal metacarpus. Scale bars = 500 µm. (E) High-magnification micrographs of articular cartilage region of the metacarpus. Scale bars = 100 µm. D = days of age, M = months of age, Y = years of age.

Histological analyses showed how the distinct metacarpal epiphyses underwent synostosis. At 3 days, the metacarpus had two distinct growth plates (*i.e.,* physes) connected by a non-ossified, non-cartilaginous, periosteum-like fibrous tissue (**Fig. 6C & D**). Between 3 days and 1.5 months of age, the two physes joined and the previous fibrous tissue exhibited robust glycosaminoglycan (GAG) deposition, which continued through the epiphyseal region. Between 1.5 and 3 months, endochondral ossification initiated in the interstitial fibrocartilaginous matrix, starting at the border with the conjoined physes. This process led to epiphyseal synostosis by 9 months of age. From 9 to 12 months old, the metacarpals continued longitudinal growth. Throughout this epiphyseal synostosis, the physes maintained a fibrocartilaginous bridge, devoid of chondrocyte columns between physes. Between 12 months and 3.5 years, the physis closed and the distal metacarpal bone remodeled to form a singular continuous network of cancellous bone.

The articular cartilage also matured over postnatal development. At 3 days, the distal portion of the metacarpal bone exhibited a coherent articular cartilage surface with robust GAG content (**Fig. 6E**). The articular cartilage matured and condensed throughout postnatal development, forming superficial, transitional, and deep zones by 3 months of age and reduced GAG content by 12 months of age.

## 4.0 Discussion

Here, we characterized the dynamics of postnatal bone development and epiphyseal synostosis in the caprine autopod from birth through adulthood. P1 bone accrual was driven by growth patterning, while cross-sectional structural morphodynamics were maintained as ambulatory loading increased. While P1 distal trabeculae increased in thickness and number, metacarpal epiphyseal trabecular bone increased primarily by thickening of pre-existing trabeculae. The two bones that fuse to form the caprine metacarpus underwent diaphyseal synostosis, prenatally, but epiphyseal synostosis postnatally, prior to growth plate closure, through fibrocartilaginous endochondral ossification. Together, these data provide new insights into precocial bone development under ambulatory loading. Our findings may guide future caprine mechanobiology studies to mechanistically understand bone development and adaptation to ambulatory loading.

P1 cortical bone development was driven by both patterned growth and ambulatory loading. Cortical bone rapidly condensed within months, followed by continuous cortical bone accrual through adulthood. Notably, the body mass-normalized polar moment of inertia (*J*/mass) was maintained over developmental time, suggesting morphodynamic adaptation of structural distribution to resist bending deformation by ambulatory loading. These findings demonstrate the early functional capacity of the precocial skeleton, which persists into adulthood. The precocial skeleton develops so robustly that it can even develop normally after preterm birth and reduced birthweight (Magrini et al. 2023). In humans, which are altricial, peripubertal physical activity increases bone formation and establishes bone structural properties that maintain bone mass and strength (Palaiothodorou and Vagenas 2024; Weatherholt and Warden 2018), which persists into adulthood (Ducher et al. 2006; Warden et al. 2014). These findings support the notion that early and peripubertal impact loading can preserve adult bone strength and reduce fracture risk (Cooper et al. 2006; Ireland et al. 2014). Future studies can use the caprine model system to determine how altered perinatal mechanical loading impacts long-term skeletal health.

Trabecular architecture matured in P1 and metacarpal bones at different rates, but both were significantly different than altricial developmental dynamics. Trabecular bone only makes up 20% of total bone mass (Ott 2018), but the trabecular microarchitecture is crucial for determining bone mechanical properties (Liu et al. 2006; Ulrich et al. 1999). The developing caprine autopod of the lower forelimb is useful for studying adaptive trabecular bone growth because precocial species experience progressive biomechanical loading from gestation, through ambulation at birth, and into adulthood. Metacarpal distal epiphyseal BV/TV increased at a slower rate than P1 distal BV/TV, likely because metacarpal trabecular bone formed by trabecular bone apposition, rather than de novo trabecula formation. Since the metacarpal bone is more proximal than the P1 bone, it develops earlier in utero (Saunders 1948; Summerbell et al. 1973); thus, the metacarpal trabecular network may have formed in utero and only developed by bone accrual postnatally. Despite different mechanisms of development, trabecular BV/TV increased continuously after birth in the P1 and metacarpal bones, which is distinct from altricial trabecular bone dynamics. Altricial species experience a reduced loading state from birth until the onset of ambulation, which may explain the loss in trabecular bone volume fraction during early postnatal development (Acquaah et al. 2015). Once humans begin walking, trabecular BV/TV continuously increases coincidently with progressive loading (Ryan and Krovitz 2006; Gosman and Ketcham 2009). These post-ambulatory BV/TV increases mirror the progressive increases in trabecular bone observed in our study. Our findings contribute to the growing body of literature describing how precocial animals adapt to the biomechanical loading of ambulation (Gorissen et al. 2016; Skedros et al. 2004; Skedros et al. 2007).

The epiphyseal synostosis of the caprine metacarpal bones is distinct from other known mechanisms of endochondral bone formation or bone fusion. Here we show that the metacarpal physes and epiphyses are initially separated by a non-cartilaginous fibrous template, which becomes chondrogenic and subsequently undergoes fibrocartilaginous endochondral ossification. This rudimentary fibrous tissue is distinct from the cartilaginous rudiments that give rise to endochondral primary bone development (Berendsen and Olsen 2015), or the cartilage template that initiates the fibrocartilaginous enthesis (Killian 2022; Blitz et al. 2013; Sugimoto et al. 2013). The epiphyseal synostosis is also distinct from the fusion of flat bones of the skull. Like the metacarpal epiphyses, the sutures of the skull feature a flexible fibrous tissue, which allows for brain growth and development. However, suture fuse by intramembranous ossification, without a cartilaginous intermediate (Opperman 2000). Synostosis of metacarpal epiphyseal bone is perhaps most analogous to endochondral bone fracture healing (Bahney et al. 2019; Einhorn and Gerstenfeld 2015). The progression of fibrous to cartilaginous to bone tissue is like fracture healing, though fracture features an acute inflammatory phase and less robust fibrous tissue formation. Biomechanical stimuli are key regulators of endochondral ossification (Kozhemyakina et al. 2015; Collins et al. 2024), suture fusion (Khonsari et al. 2013; Sanchez-Lara et al. 2010), enthesis development (Killian 2022), and fracture healing (Goodship and Kenwright 1985; Matziolis et al. 2006; Herberg et al. 2019; McDermott et al. 2019; Kegelman et al. 2021) and may play a role in metacarpal epiphyseal synostosis. Interrogating the mechanical basis of metacarpal epiphyseal synostosis could lead to novel insights into the underlying mechanisms.

Together with a growing body of literature on precocial bone formation, these development dynamics data may form a foundation on which to uncover how mechanical forces regulate skeletal development, disease, and regeneration. One limitation of this study is that we could not account for sex differences. To enable sufficient numbers and ages, given housing and breeding constraints (Burmeister et al. 2022), we used male and female goats from the Translational Research Unit (TRU) goat herd at University College Dublin Lyons Research Farm. This led to different numbers of male and female goats in each group. Despite this limitation, our data exhibit high coefficients of determination (R^2^). This suggests that our methods were sufficiently robust to capture the dynamics of postnatal bone development. Our ability to get consistent results is in agreement with the benefits of using genetically diverse populations in bone research (Migotsky et al. 2024). In future studies, we aim to directly interrogate sex as a biological variable among animals of more homogenous backgrounds (Nelson and Megyesi 2004).

## 5.0 Acknowledgements

This work was supported by the European Union’s Horizon 2020 Research and Innovation Programme [EURONANOMED2017-77 to PB], Research Ireland [21/FFP-A/9090 to NCN], the Science Foundation Ireland [SFI/16/ENM-ERA/3458 to PB], the National Institute of Arthritis, Musculoskeletal and Skin Diseases (NIAMS) of the National Institutes of Health (NIH) [R01-AR074948 & R01-AR073809 to JDB], Penn Center for Musculoskeletal Disorders [NIH/NIAMS P30-AR069619 to JDB], the Center for Engineering Mechanobiology (CEMB) [NSF CMMI: 15-48571 to JDB], the University of Pennsylvania Institutional Research and Academic Career Development Award (IRACDA) [NIH/NIGMS K12GM081259 to CJP], and the University of Pennsylvania Center for Undergraduate Research and Fellowships [CURF to CJP and ME]. We thank the University of Pennsylvania University Archives and Records Center for providing access to the original plates and reproductions of the photographs of Eadweard Muybridge (**Fig. 1**), whose photographic laboratory would be visible from JDB’s office window.

## 6.0 Author contributions

Concept/design: CJP, MCL, PAJB, JDB, NCN. Acquisition of data: CJP, ME, EB, AW. Data analysis/interpretation: CJP, ME, EB, AW, PAJB, JDB, NCN. Drafting of the manuscript: CJP, PAJB, JDB, NCN. Critical revision of the manuscript: CJP, ME, EB, AW, KV, MCL, PAJB, JDB, NCN. Approval of the article: CJP, ME, EB, AW, KV, MCL, PAJB, JDB, NCN.

